# Modulating Prestimulus Alpha and Beta Power with tRNS Establishes Their Causal Role in Visual Perception

**DOI:** 10.1101/2024.12.11.627917

**Authors:** Jinwen Wei, Huiru Zou, Qianyuan Tang, Ziqing Yao, Gan Huang, Zhen Liang, Li Zhang, Changsong Zhou, Zhiguo Zhang

## Abstract

Variability in visual perception in response to consistent stimuli is a fundamental phenomenon linked to fluctuations in prestimulus low-frequency neural oscillations—particularly in the alpha (8–13 Hz) and beta (13–30 Hz) bands—typically measured by their power in electroencephalography (EEG) signals. However, the causal role of these prestimulus alpha and beta power fluctuations in visual perception remains unestablished. In this study, we investigated whether prestimulus alpha and beta power causally affect visual perception using transcranial random noise stimulation (tRNS). In a sham-controlled, single-blind, within-subject design, 29 participants performed a visual detection task while receiving occipital tRNS. Online functional near-infrared spectroscopy (fNIRS) was used to measure cortical excitability during stimulation, and offline EEG signals were collected after stimulation. Mental fatigue was incorporated as a state-dependent factor influencing tRNS effects. Our findings demonstrate that, primarily under low fatigue states, tRNS increased cortical excitability during stimulation (indicated by increased fNIRS oxyhemoglobin amplitude), decreased subsequent prestimulus EEG alpha and beta power, and consequently reduced the visual contrast threshold (VCT), indicating enhanced visual perception. Sensitivity analysis revealed that alpha oscillations contributed more significantly to visual perception than beta oscillations under low fatigue. Additionally, the state-dependent effects of tRNS may result from different sensitivities of VCT to neural oscillations across fatigue states. These results provide causal evidence linking prestimulus alpha and beta power to visual perception and underscore the importance of considering brain states in neuromodulation research. Our study advances the understanding of the neural mechanisms underlying visual perception and suggests potential therapeutic applications targeting neural oscillations.

**Significance statement:** Understanding why we perceive identical visual stimuli differently is a fundamental question in neuroscience. This study provides causal evidence that prestimulus alpha and beta neural oscillations directly influence visual perception, particularly under low mental fatigue state. By using tRNS alongside fNIRS and EEG recordings, we demonstrate that modulating neural excitability can alter perceptual outcomes. Our findings highlight the importance of considering brain state—such as fatigue levels—in neuromodulation research. This work advances our understanding of the neural mechanisms underlying visual perception and opens avenues for developing targeted interventions to enhance sensory processing and cognitive functions, potentially benefiting individuals with perceptual or attentional disorders.

## Introduction

Variability in visual perception in response to consistent stimuli remains a significant topic of interest in neuroscience. This phenomenon is exemplified by an observer’s fluctuating ability to detect near-threshold visual contrast stimuli across different trials. Previous research has demonstrated that these perceptual inconsistencies are linked to prestimulus (or spontaneous) low-frequency neural oscillations, particularly within the alpha (8–13 Hz) and beta (13–30 Hz) ranges preceding stimulus onset (1). An inverse relationship between the amplitude of these prestimulus low-frequency oscillations and the ability to detect near-threshold visual stimuli has been extensively established, suggesting that suppression of alpha or beta activity in the posterior cortex may enhance the baseline sensitivity of sensory systems (2–12).

However, despite the temporal precedence of these neural oscillations in visual perception, current studies have primarily provided correlational rather than causal evidence for their link (1, 8). In other words, it remains unknown whether prestimulus alpha and beta oscillations causally affect visual perception. This gap in knowledge is due to the predominant use of non-invasive neural imaging techniques, such as electroencephalography (EEG) and functional magnetic resonance imaging (fMRI), which primarily establish correlational relationships between neural features and behaviours (13). The neuroscience community has emphasized the necessity of establishing causal relationships between physiological features (e.g., neural oscillations) and cognitive behaviours (e.g., visual perception) (14, 15). Establishing causality would significantly enhance our understanding of neural mechanisms and has important implications for rehabilitation and therapeutic interventions (16).

Transcranial electrical stimulation (tES), a non-invasive neuromodulation technique, has significantly advanced our exploration of causal relationships between neural dynamics and cognitive functions (17). Transcranial random noise stimulation (tRNS), in particular, has been shown to enhance cortical excitability (18), and improve visual perception (19), among other functions. By boosting cortical excitability—which related closely to EEG alpha (11, 20) and beta (21, 22) power—tRNS offers a promising approach to studying the causal effects of prestimulus alpha and beta power on visual perception.

However, two challenges remain when using tRNS in practical applications. First, recent research has shown that the efficacy of tES is influenced by specific neural states such as arousal (23), sleepiness (24), and ongoing oscillations (25). This state-dependent effect of tES is believed to contribute to the variability in modulation outcomes across studies (26). Therefore, it is crucial to consider the influence of brain states during tRNS application when establishing a causal link between prestimulus alpha/beta power and visual perception. Second, studying the online effects of tES on neural oscillations is challenging due to the contamination of EEG recordings by tES artifacts (27). Hemodynamic responses measured by fMRI or functional near-infrared spectroscopy (fNIRS) are less susceptible to these artifacts (13). Hemodynamic changes are closely linked to neural activity through neurovascular coupling, where increases in cortical excitability correlate with enhanced hemodynamic responses (28). Consequently, integrating fNIRS—which is more portable and cost-effective than fMRI—with EEG allows for a convenient and comprehensive examination of both the online hemodynamic effects and offline neural oscillations during tRNS application, overcoming the limitations posed by tES artifacts in EEG alone.

We hypothesized that tRNS would modulate both online hemodynamic responses measured by fNIRS and offline prestimulus alpha and beta power measured by EEG, thereby influencing visual perception in a state-dependent manner. Specifically, we aimed to establish a causal relationship between prestimulus alpha/beta oscillations and visual perception by demonstrating that tRNS-induced changes in these oscillations lead to altered perceptual performance, depending on the mental fatigue state.

In this study, we utilized a tRNS-based experimental framework, integrating fNIRS and EEG with a visual detection task in a sham-controlled, single-blind, within-subject design. By considering mental fatigue as a state variable, we addressed the state-dependent effects of tRNS. Our findings indicate that, mainly under low fatigue, occipital tRNS increased online fNIRS HbO amplitude, decreased subsequent prestimulus EEG alpha and beta power, and consequently reduced the visual contrast threshold (VCT), suggesting a causal relationship between prestimulus alpha/beta power and visual perception (see a schematic summary in Fig. 1). Further sensitivity analysis showed that alpha oscillations contribute more to visual perception than beta oscillations under low fatigue, and that state-dependent tRNS effects may result from different sensitivities to VCT between fatigue states.

**Fig. 1.**
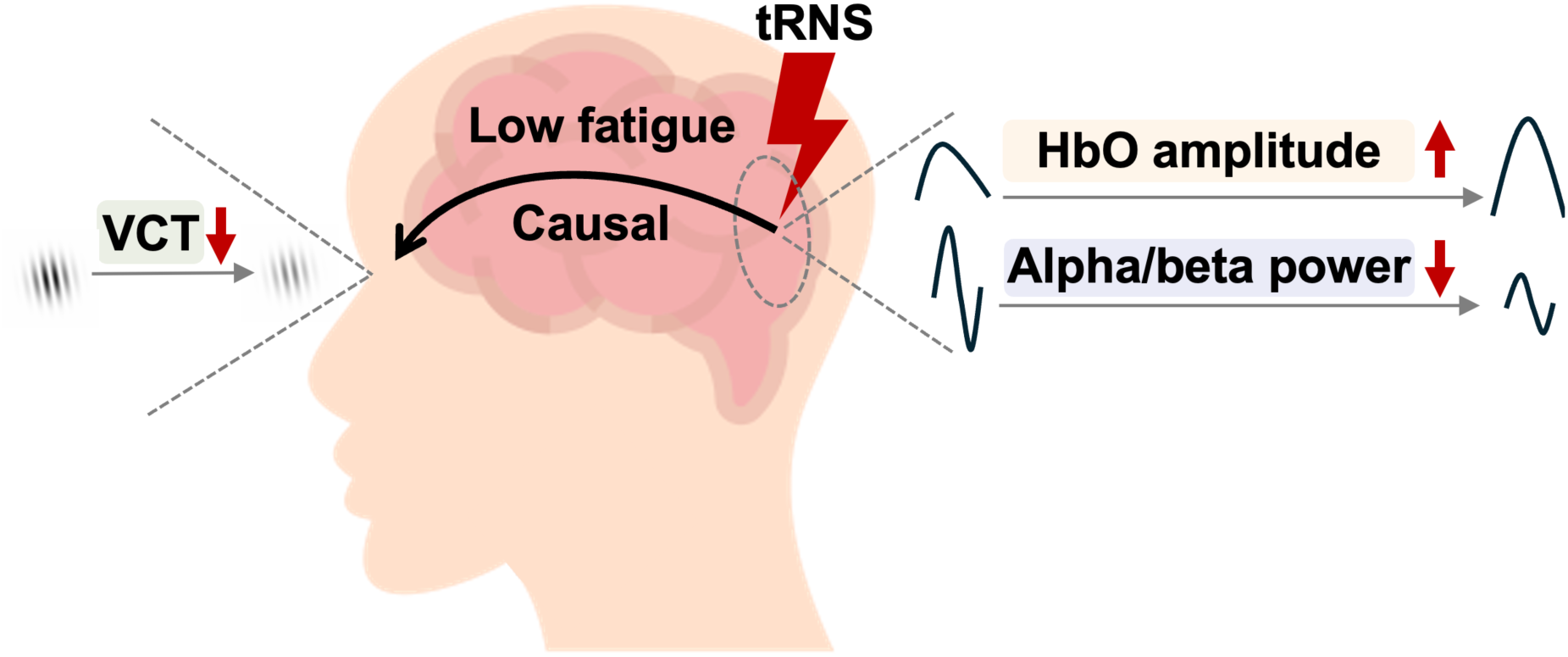
A schematic summary of our key findings. Under low fatigue, occipital tRNS increased online fNIRS HbO amplitude, decreased subsequent prestimulus EEG alpha and beta power, and consequently reduced the VCT, suggesting a causal relationship between prestimulus alpha/beta power and visual perception.

## Results

### tRNS Affects HbO Amplitude Without Considering Fatigue

To evaluate the potential causal role of prestimulus alpha and beta power in visual perception, we conducted a visual detection task combined with fNIRS/EEG recording and tRNS in 29 healthy participants (15 females; mean age, 22.7 ± 1.9 years). Participants performed a visual detection task where Gabor patches with near-threshold contrasts were presented with a probability of 60% (Fig. 2A). The experiment consisted of five blocks, each separated by a 5-minute rest interval. Each block comprised 80 trials, resulting in approximately 48 trials with stimulus presentation per block (80 × 60%). Visual contrast threshold (VCT) was estimated every 40 trials using a cumulative Gaussian function following our previous study (29). Each participant completed the task twice—once with tRNS and once with sham stimulation—with an interval of 3–7 days between sessions.

**Fig. 2.**
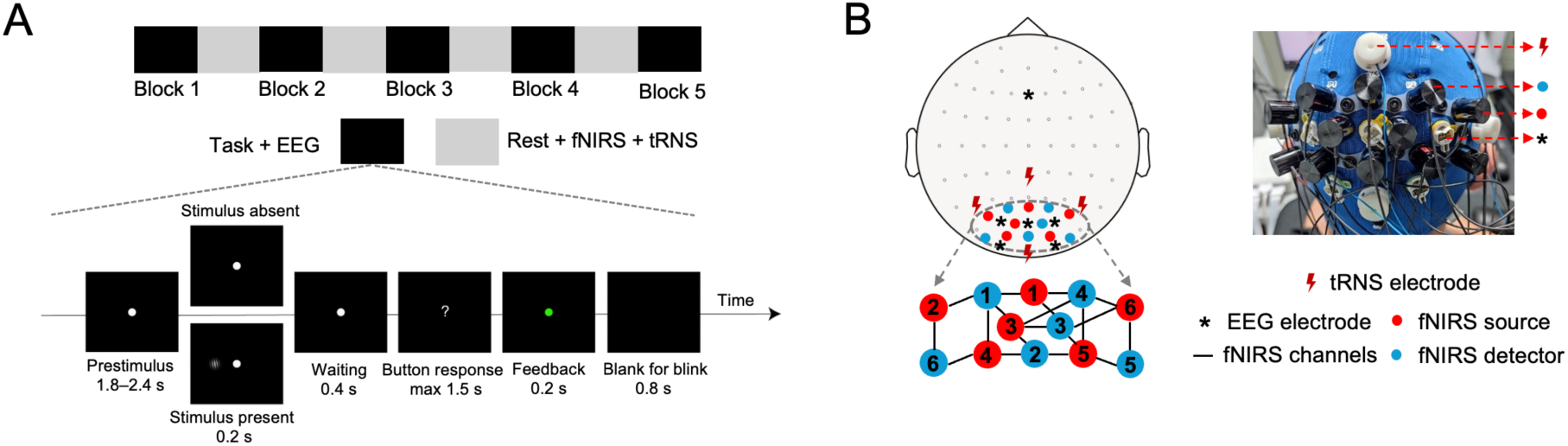
Experimental setup. (*A*) The experiment consisted of five blocks, each separated by a 5-minute resting-state period during which tRNS was applied, and fNIRS signals were recorded. Each block contained 80 trials. In 60% of trials, participants were presented with a near-threshold visual stimulus (a Gabor patch) for 0.2 s on either the left or right side of a central fixation dot (50% probability for each side). After a 0.4 s interval, the fixation dot changed to a question mark, prompting participants to indicate whether they perceived the stimulus by pressing a button. Feedback was provided via a green (correct) or red (incorrect) dot for 0.2 s, followed by a blank screen for blinking. The next trial began after a variable interval of 1.8–2.4 s. (*B*) EEG signals were recorded from occipital electrodes: O1, O2, PO3, POz, and PO4, with FCz as the reference electrode. Stimulation electrodes were placed over CPz, P5, P6, and Oz. Two stimulation conditions—sham and tRNS—were implemented in a counterbalanced, single-blind, within-subject design on two different days. fNIRS sources and detectors were positioned between tRNS and EEG electrodes (see Methods for details).

Fatigue ratings were collected using a 7-point scale before and after each block, with the mean score representing the prevalent fatigue state for each block. EEG data were collected during task blocks, while tRNS with a high-frequency band of 100–640 Hz was applied and fNIRS data were collected over the occipital lobe during rest periods (Fig. 2B). It should be noted that fNIRS data were collected during the tRNS implementation, while EEG data were collected afterward (Fig. 2A).

We first analysed the effects of tRNS on HbO amplitude, prestimulus alpha and beta power, and VCT without considering fatigue states, using Bayesian linear mixed models (see details in Methods). Evidence supporting a hypothesis was considered compelling if the posterior probability (*Pr*) exceeded 97.5% and the 95% highest posterior density (*HPD*) interval did not include zero. Compared to the sham condition, tRNS increased HbO amplitude only in the 5^th^ block (Fig. 3A; *E* (*μ*_Sham-tRNS, Block 5_) = −0.169, *HPD* = [−0.278, −0.063], *Pr* (*μ*_Sham-tRNS, Block 5_<0) = 0.999; see complete statistics in Table S1). However, tRNS did not modulate prestimulus alpha power (Fig. 3B; all *Pr*s < 0.975; see Table S2), prestimulus beta power (Fig. 3C; all *Pr*s < 0.975; see Table S3), or VCT (Fig. 3D; all *Pr*s < 0.975; see Table S4).

**Fig. 3.**
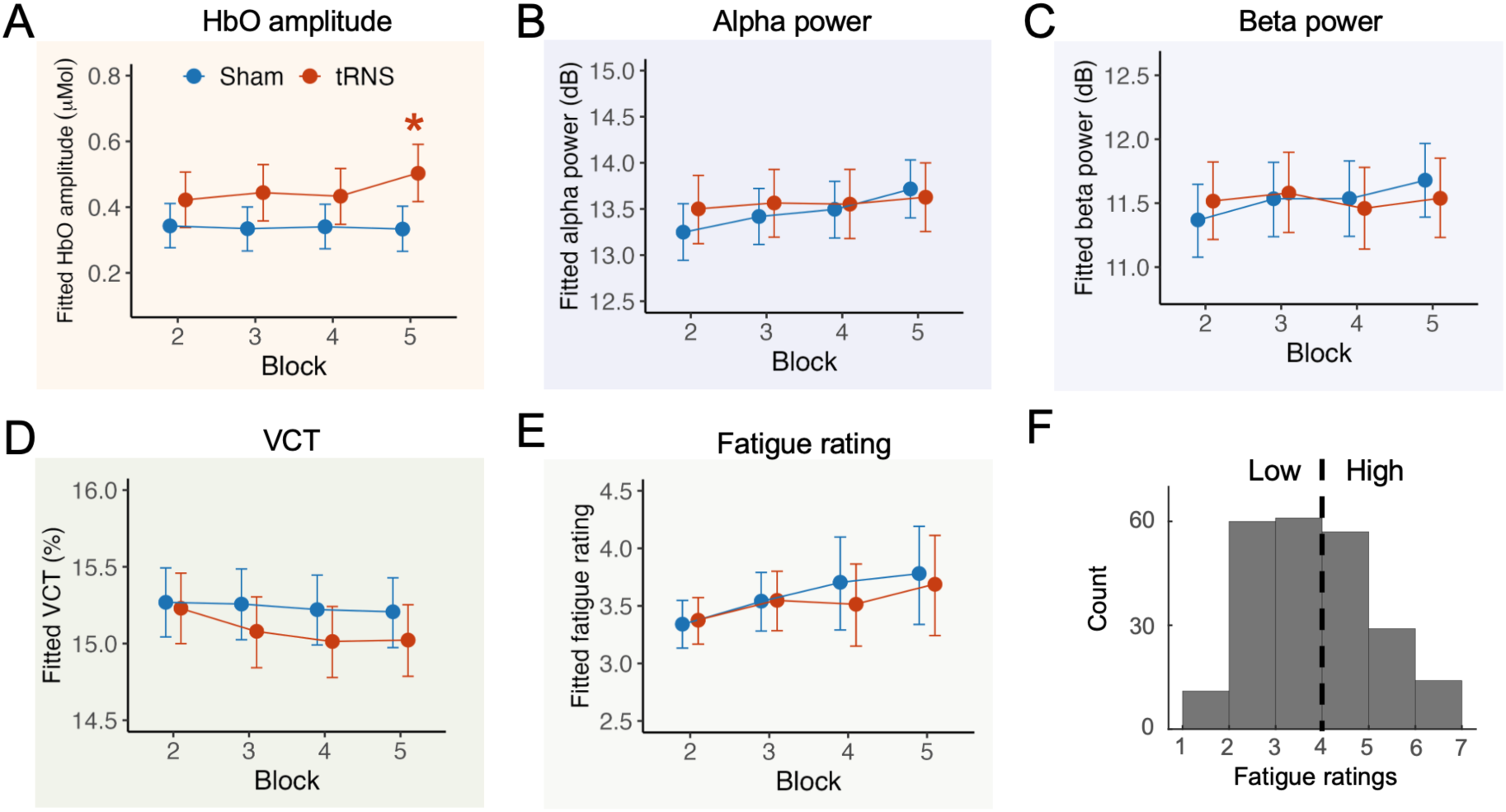
tRNS effects without considering fatigue. (*A*) tRNS increased HbO amplitude in the fifth block compared to sham stimulation. The asterisk (*) indicates a posterior probability exceeding 97.5%, with a 95% HPD interval that does not include zero, providing compelling evidence. (*B*) tRNS did not affect prestimulus alpha power. (*C*) tRNS did not affect prestimulus beta power. (*D*) tRNS did not affect VCT. (*E*) Fatigue ratings were unaffected by tRNS compared to sham stimulation, confirming fatigue as an independent variable for categorization. (*F*) Given the independence of fatigue ratings from stimulation conditions, fatigue ratings were categorized into two groups: low fatigue (< 4) and high fatigue (≥ 4).

Considering accumulating evidence that brain states influence the effects of tES (26), we included fatigue as an interaction factor in subsequent analyses. Before examining state-dependent effects, we verified that fatigue ratings were not influenced by tRNS. The results confirmed that tRNS did not affect fatigue ratings compared to the sham condition (Fig. 3E; all *Pr*s < 0.975; see Table S5), affirming the independence of fatigue from the stimulation conditions. Consequently, data samples—including fNIRS HbO amplitude, EEG power in different frequency bands, and VCT—were categorized into low and high fatigue groups using a threshold of 4 on the 7-point scale (Fig. 3F), distinguishing low fatigue (< 4) from high fatigue (≥ 4).

### tRNS Increased HbO amplitude, Decreased Prestimulus Alpha/Beta Power, and Consequently Reduced VCT Under Low Fatigue

We next analysed the effects of tRNS while considering fatigue as an interaction factor, using Bayesian linear mixed models. In the 5^th^ block under low fatigue, compared to the sham condition, tRNS notably increased HbO amplitude (Fig. 4A; *E* (*μ*_Sham-tRNS_) = −0.276, *HPD* = [−0.405, −0.137], *Pr* (*μ*_Sham-tRNS_ < 0) = 1; see Table S6), decreased prestimulus alpha power (Fig. 4B; *E* (*μ*_Sham-tRNS_) = 0.582, *HPD* = [0.034, 1.162], *Pr* (*μ*_Sham-tRNS_ > 0) = 0.977; see Table S7), decreased prestimulus beta power (Fig. 4C; *E* (*μ*_Sham-tRNS_) = 0.482, *HPD* = [0.047, 0.927], *Pr* (*μ*_Sham-tRNS_ > 0) = 0.983; see Table S8), and reduced VCT (Fig. 4D; *E* (*μ*_Sham-tRNS_) = 0.307, *HPD* = [0.001, 0.603], *Pr* (*μ*_Sham-tRNS_ > 0) = 0.978; see Table S9). In contrast, tRNS did not have significant effects in other blocks or under high fatigue (Fig. 4).

**Fig. 4.**
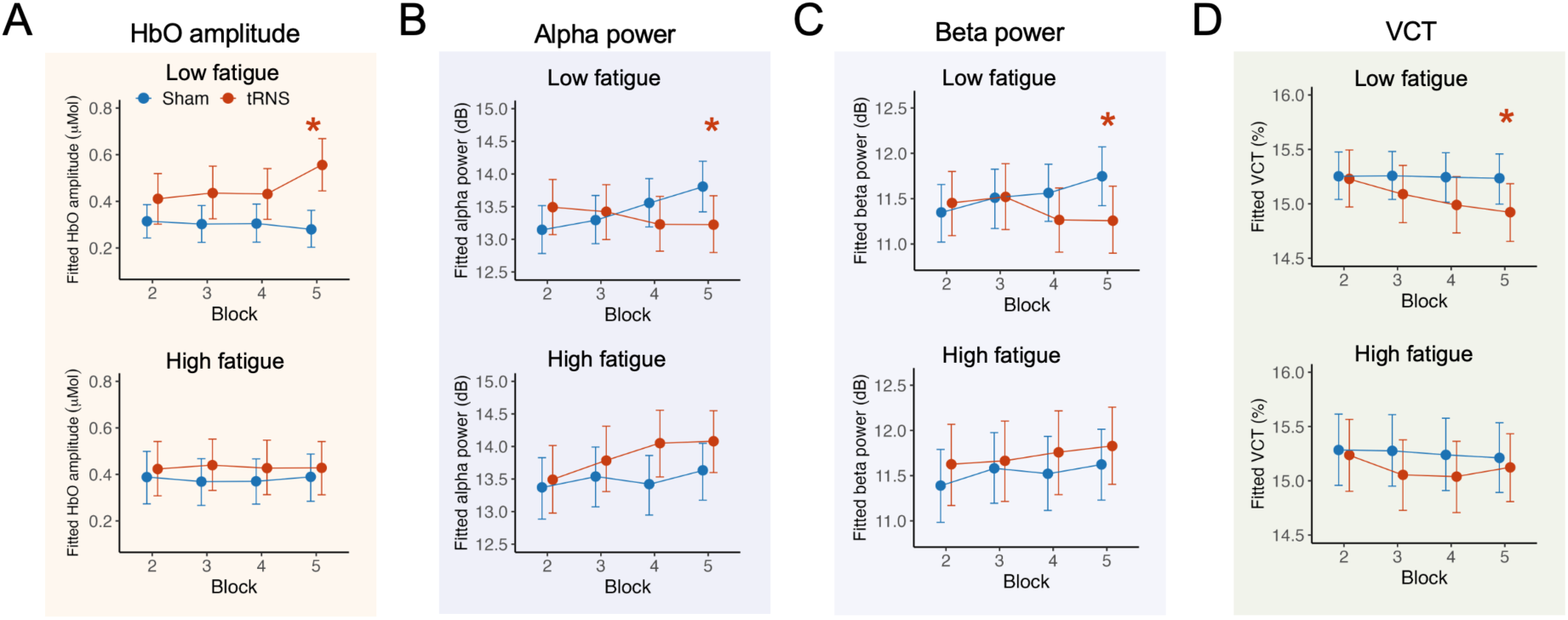
State-dependent effects of tRNS on fNIRS HbO amplitude, prestimulus EEG alpha and beta power, and VCT. (*A*) Under low fatigue, tRNS increased fNIRS HbO amplitude in Block 5 compared to sham stimulation. Under high fatigue, tRNS did not influence fNIRS HbO amplitude in any block. The asterisk (*) indicates a posterior probability (*Pr*) exceeding 97.5%, with a 95% *HPD* interval that does not include zero, providing compelling evidence. (*B*) Under low fatigue, tRNS decreased prestimulus alpha power in Block 5 compared to sham stimulation. Under high fatigue, tRNS did not influence prestimulus alpha power in any block. (*C*) Under low fatigue, tRNS decreased prestimulus beta power in Block 5 compared to sham stimulation. Under high fatigue, tRNS did not influence prestimulus beta power in any block. (*D*) Under low fatigue, tRNS decreased VCT in Block 5 compared to sham stimulation. Under high fatigue, tRNS did not influence VCT in any block.

Given the direct physiological impact of tES (17, 30, 31), it is plausible that tRNS directly affected HbO amplitude during stimulation, subsequently influencing prestimulus alpha and beta power, and consequently leading to the observed decrease in VCT under low fatigue in the same block.

### tRNS Did Not Affect Prestimulus Delta, Theta, or Gamma Power

To ensure that the observed variations in VCT were not attributable to alterations in prestimulus power in other frequency bands, we investigated how tRNS affected prestimulus power in the delta (0.5−4 Hz), theta (4−8 Hz), and gamma (30−45 Hz) bands. The results showed that, compared to the sham condition, tRNS did not affect prestimulus power in these frequency bands under low or high fatigue states (Fig. 5; all *Pr*s < 0.975; see complete statistics in Table S10−12). Therefore, tRNS did not influence VCT through changes in delta, theta, and gamma band power.

**Fig. 5.**
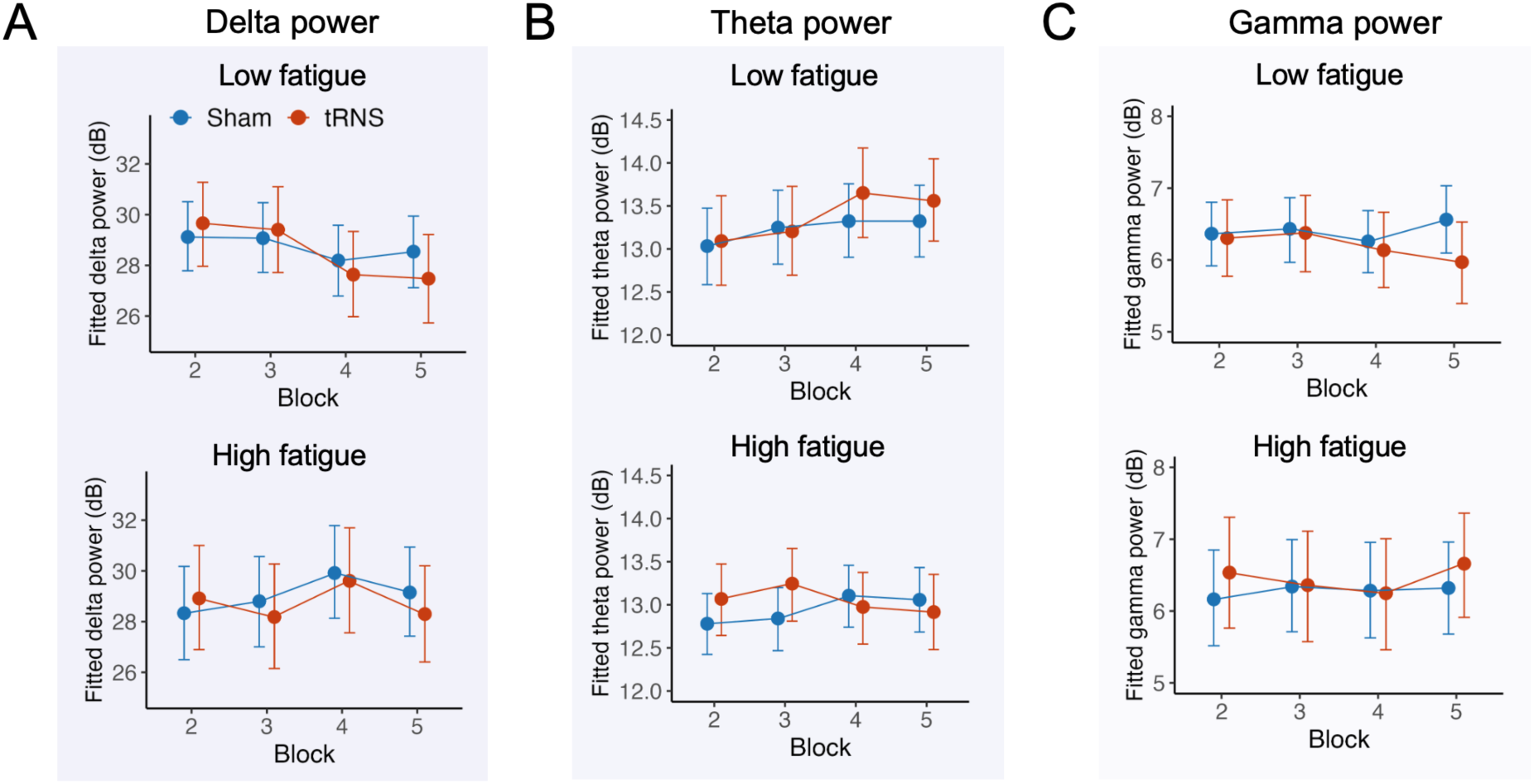
Effects of tRNS on prestimulus delta, theta, and gamma power. (*A*) tRNS did not affect prestimulus delta power under either low or high fatigue states. (*B*) tRNS did not affect prestimulus theta power under either low or high fatigue states. (*C*) tRNS did not affect prestimulus gamma power either both low or high fatigue states.

### VCT is more sensitive to alpha power than beta power under low fatigue

The above findings indicated that both alpha and beta band powers might causally affect VCT primarily under low fatigue conditions. To examine the respective contributions of alpha and beta powers to VCT and to understand the underlying mechanisms of tRNS effects under low fatigue, we conducted a sensitivity analysis of different EEG frequency band powers on VCT under low and high fatigue states.

Specifically, we trained two neural network models to capture potential nonlinear relationships between EEG band power and VCT in the sham group under low and high fatigue conditions (Fig. 6A). Both models had identical architectures and hyperparameters. The training data comprised single-trial EEG power features and corresponding VCT measurements from the sham group, with 6,640 samples for low fatigue and 4,960 samples for high fatigue. We focused on the sham group to establish baseline effects without tRNS influence. Each sample included 25 input features representing power across 5 frequency bands and 5 electrodes, and the output was the VCT.

**Fig. 6.**
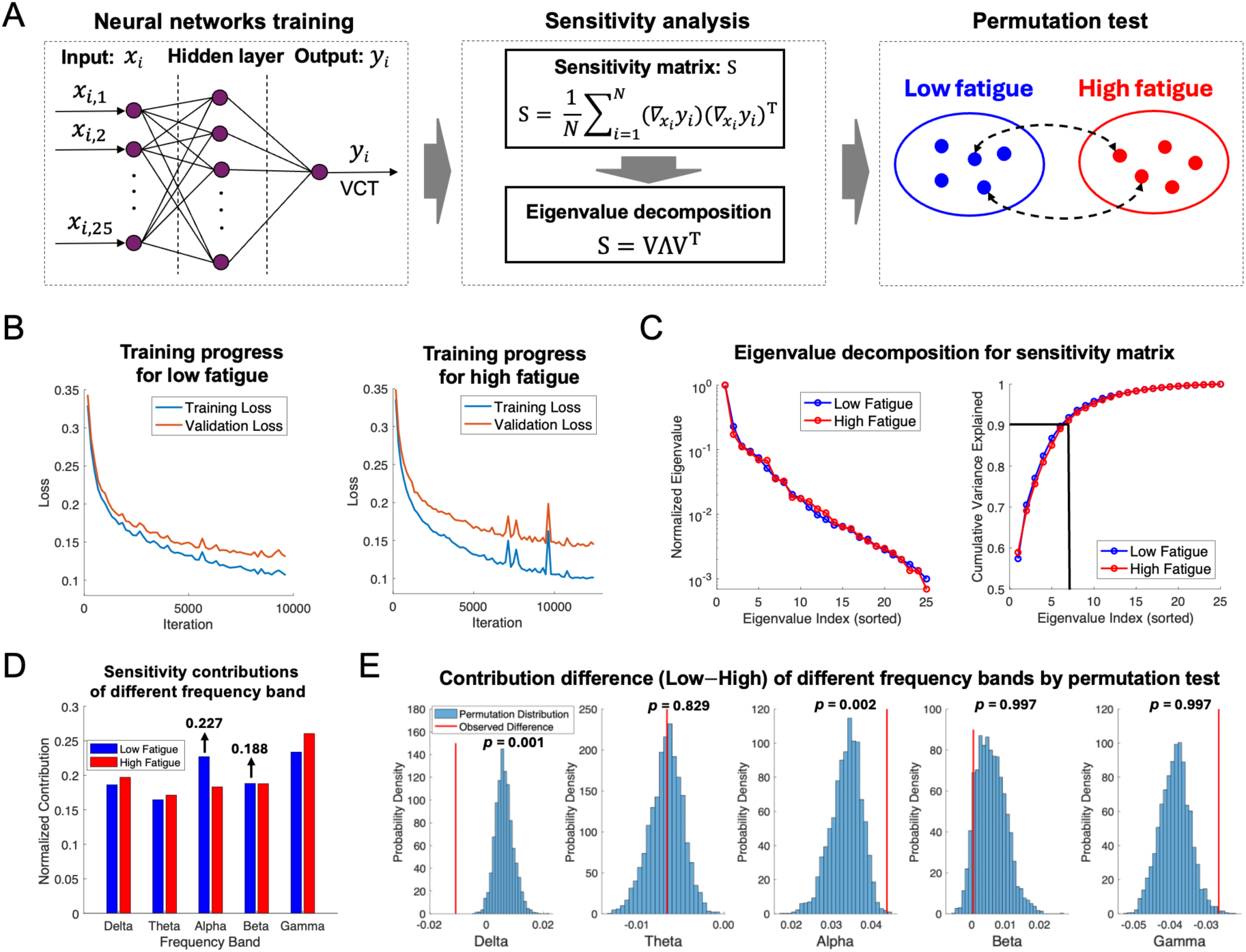
Sensitivity analysis of EEG frequency bands on VCT in the sham group. (*A*) Two neural network models were trained to predict VCT using single-trial EEG power across different frequency bands under low and high fatigue states in the sham group. After training, sensitivity analysis was performed by computing the sensitivity matrix and conducting eigenvalue decomposition. Permutation tests were used to compare the sensitivity contributions of different frequency bands between low and high fatigue states (see Methods for details). (*B*) To prevent overfitting, samples under each fatigue state were divided into training and validation subsets. Early stopping resulted in well-generalized models for both low and high fatigue conditions, with validation loss slightly lower than training loss, indicating minimal overfitting. The similar loss values across conditions suggest the models are comparable for analysis. (*C*) Eigenvalue decomposition of the sensitivity matrix was performed. Eigenvalues explaining 90% of the variance were selected to calculate the sensitivity contributions of different frequency bands to VCT. (*D*) Under low fatigue, the alpha band showed a larger sensitivity contribution to VCT than the beta band. (*E*) The alpha band had higher sensitivity under low fatigue compared to high fatigue, whereas the beta band did not show a significant difference between fatigue states.

To prevent overfitting, we divided each dataset into training and validation subsets in an 8:2 ratio and employed early stopping during training (Fig. 6B). After training the models, we performed sensitivity analysis by computing the sensitivity matrix and conducting eigenvalue decomposition (Fig. 6A, middle panel). We selected eigenvalues that explained 90% of the variance to calculate the sensitivity contributions of each frequency band to VCT (Fig. 6C). The results showed that the contribution of the alpha band (0.227) was greater than that of the beta band (0.188), suggesting that alpha power has a more significant impact on VCT than beta power under low fatigue (Fig. 6D).

To explore potential reasons why tRNS effects were observed only under low fatigue, we conducted permutation tests to compare the sensitivity contributions of different frequency bands between low and high fatigue states. We randomly permuted all samples 10,000 times and calculated the contribution of each frequency band in each permutation using the same method as above. As shown in Fig. 6E, the delta band had a higher contribution under high fatigue (*p*_corrected_ = 0.001), while the alpha band had higher sensitivity under low fatigue (*p*_corrected_ = 0.002). In contrast, there were no significant differences in the contributions of the theta (*p*_corrected_ = 0.829), beta (*p*_corrected_ = 0.997), and gamma (*p*_corrected_ = 0.997) bands between low and high fatigue states. These findings imply that the lack of tRNS effects on alpha and beta bands under high fatigue may be due to their lower sensitivity in influencing VCT. The justification for the hyperparameters we used can be found in Table S13 and Fig. S2.

## Discussion

This study aimed to clarify the causal relationship between prestimulus low-frequency oscillation and visual perception. Our results demonstrate that tRNS, particularly under brain state of low fatigue, led to increased online fNIRS HbO amplitude, decreased subsequent prestimulus EEG alpha and beta power, and a reduction in VCT. These findings support a causal link between prestimulus alpha/beta power and visual perception. Furthermore, sensitivity analyses revealed that alpha oscillations have a more significant influence on visual perception than beta oscillations under low fatigue, suggesting that the state-dependent effects of tRNS may stem from varying sensitivities of VCT to neural oscillations across different fatigue states.

We first analysed the effects of tRNS without considering fatigue states. The stimulation effects were minimal compared to the state-dependent effects shown in Figures 4 and 5. This contrast, attributable to fatigue states—or brain states in a broader sense—may be one of the main reasons for inconsistent results in existing tES studies (26, 32, 33, 34). This observation aligns with the trend that an increasing number of tES studies are incorporating brain state factors into the analysis of stimulation effects (26, 35, 36). We then analysed the effects of tRNS on fatigue states and found that fatigue was not modulated by tRNS. These null effects suggest that mental fatigue, which commonly correlates with arousal (37–39), may involve different mechanisms from those underlying alpha oscillations, which showed state-dependent effects. Similar dissociations have been found, indicating that arousal and alpha oscillations may have interconnected but non-overlapping roles in visual perception (40). After determining the independence of mental fatigue from stimulation effects, we classified the data samples using a fixed threshold of 4 on the fatigue scale, following the same approach as in our previous study (29). Potential bias due to this binary classification can be alleviated by two points: first, the low and high fatigue samples were relatively comparable in number (low: 132; high: 100); second, the Bayesian linear mixed model is capable of handling sample imbalance, thus providing robust results (41, 42).

The state-dependent effects of tRNS suggest a causal role of prestimulus alpha and beta oscillation power in visual perception. This establishment of causality satisfies two practical requirements for a causal factor (43): first, existence of temporal precedence, demonstrated by the “prestimulus” neural features; second, changes from intervention, through the modulation of neural oscillations by tRNS in this study. The possibility that other frequency bands might have effects was ruled out by analysing tRNS effects on delta, theta, and gamma oscillation power, as shown in Fig. 5. While numerous studies have shown correlations between prestimulus alpha/beta oscillations and visual perception (1), our study provides causal evidence, underscoring significant implications for the neural mechanisms of visual perception and related therapeutic applications.

Our findings have significant implications for developing interventions aimed at enhancing sensory processing and cognitive functions, particularly for individuals with perceptual or attentional disorders. Disorders such as amblyopia, attention deficit-hyperactivity disorder (ADHD), and visual neglect are associated with abnormalities in neural oscillations and perceptual deficits (44, 45). By demonstrating that modulating prestimulus alpha and beta power can causally influence visual perception, our study suggests that tRNS could serve as a non-invasive neuromodulation technique for therapeutic interventions. tRNS has been shown to enhance cortical excitability and improve cognitive functions without discomfort (19). Previous research indicates that tRNS can facilitate perceptual learning in amblyopia patients (46) and improve attentional performance in individuals with ADHD (47, 48), and etc. Considering the state-dependent nature of tRNS effects, personalized interventions that account for individual brain states, such as fatigue levels, could optimize efficacy (26). Future studies should explore the clinical applications of tRNS in these populations, potentially leading to novel treatments for perceptual and attentional disorders.

It should be noted that the prestimulus alpha and beta oscillations we investigated were “online” with visual perceptual behaviour (VCT) but “offline” with respect to tRNS application. The offline tRNS effects on neural oscillations can be traced back to the neural excitability induced by tRNS, as demonstrated by increased fNIRS HbO amplitude. Therefore, our results indicate a time-lagged relationship between neurovascular responses and neural oscillatory changes. Moreover, both the online and offline effects exhibited an accumulation effect; that is, tRNS effects became noticeable after several blocks of stimulation. Similar accumulation effects were observed in our previous study (29) and can be echoed by studies showing that incremental changes due to neural stimulation may accumulate over time, leading to a threshold-crossing effect (49).

Furthermore, the two potential causal factors, prestimulus alpha and beta power, may contribute differently to visual perception, and the causal effects were absent under high fatigue. To address these issues, we performed sensitivity analysis. The results showed that under low fatigue, alpha oscillations contributed more than beta oscillations to visual perception. This finding is consistent with the dominance of alpha oscillations reported in previous studies (1). Additionally, alpha oscillations had greater sensitivity under low fatigue than high fatigue, which may explain the lack of tRNS effects under high fatigue. Considering the higher sensitivity of other bands, such as delta, under high fatigue (as shown in Fig. 6), it is reasonable to deduce that visual perception under different fatigue states is dominated by different frequency bands. Since tRNS mainly affects neural excitability (18, 19), manifested through alpha and beta oscillations (11, 22), and more sensitive oscillations under high fatigue, such as delta, were not influenced by tRNS, this could explain the absence of tRNS effects under high fatigue.

There are some limitations in our study. First, while subjective ratings provided a “gold standard” of mental fatigue, they are still relatively subjective compared to some objective physiological measures, such as pupil size (38). Therefore, future work could employ such measures to quantify brain states when studying state-dependent effects of tES. Second, we mainly collected and analysed localized neural activity driven by our hypothesis. It would be beneficial to also analyse neural activity in other brain areas, which could provide a more comprehensive understanding of the neural mechanisms involved.

In conclusion, our study supports the causal relationship between prestimulus alpha and beta power and visual perception by combining fNIRS/EEG recordings and tRNS. The establishment of this causal link not only advances our understanding of the neural mechanisms underlying visual perception but also has broader implications for developing interventions aimed at enhancing sensory processing and cognitive functions.

## Methods

### Participants

This study included 29 participants (15 females; mean age, 22.7 ± 1.9 years) after excluding nine individuals due to misunderstanding of instructions, excessive EEG or fNIRS artifacts, or discomfort during stimulation. The original sample size of 38 was consistent with our previous studies (12, 29) to yield more comparable results. All participants were right-handed, on-campus students with normal or corrected-to-normal vision. Exclusion criteria were stringent, encompassing left-handedness; a history of seizures or head injuries; current use of psychotropic medication; presence of metal implants in the head; implanted electronic devices; psychiatric disorders; tinnitus; or participation in other neuromodulation experiments within the last three months. Prior to the experiment, participants were instructed to rest adequately and abstain from caffeine or alcohol. Informed consent was obtained from all participants. The study was approved by the local ethics committee at Shenzhen University.

### Experimental Design

We employed a visual detection task adapted from previous studies (8, 9), implemented in MATLAB (MathWorks, Natick, MA) using Psychophysics Toolbox 3. The experiment was conducted in a dark, acoustically isolated chamber. Visual stimuli consisted of near-threshold Gabor patches displayed on a 21-inch LCD monitor with a 100 Hz refresh rate, positioned 60 cm from the participants. These patches, tilted at 10 degrees and subtending a visual angle of 0.75 degrees, appeared on either side of a central fixation dot for 0.2 s in 60% of the trials. Participants responded by pressing buttons after a 0.4 s delay when the fixation dot changed to a question mark. Feedback was provided by displaying a green (correct) or red (incorrect) dot for 0.2 s. Each trial was followed by a blank screen interval for blinking and an inter-stimulus interval (ISI) of 1.8–2.4 s (Fig. 2A).

The experiment comprised five blocks, each separated by a 5-minute resting-state period during which tRNS was applied and fNIRS signals were recorded. EEG data were collected during the task blocks following stimulation (Fig. 2A). Block 1 served as the baseline for subsequent statistical analyses. Each block included 80 trials, with stimulus contrast adjusted dynamically to maintain a 50% hit rate using the adaptive staircase procedure QUEST (50). The QUEST algorithm was run continuously throughout the experiment, updating its priors based on the results from the preceding block (Fig. S1A). We fitted the stimulus contrast values every 40 trials within each block using a cumulative Gaussian function and estimated the VCT as the mean of this function (29).

### Transcranial Random Noise Stimulation

Transcranial random noise stimulation (tRNS) was administered using a high-definition transcranial electrical stimulator (Soterix Medical) with an M×N 33-channel configuration. Four 12-mm-diameter Ag/AgCl ring electrodes filled with conductive gel were placed over CPz, P5, P6, and Oz (Fig. 2B). The tRNS stimulation frequencies ranged from 100 to 640 Hz in the high-frequency band (19), with a maximum current intensity of 1 mA (99% of amplitude values between –0.5 and 0.5 mA), as shown in Fig. S1B.

The study followed a sham-controlled, single-blind, within-subject design with two stimulation conditions: sham and tRNS. Each participant underwent both conditions, with a 3–7 day interval between sessions. All stimulation conditions included a 10-second fade-in/fade-out period. The sham stimulation lasted only for the first 30 seconds, while tRNS was applied during the resting periods between blocks (Fig. 2A).

Participants provided self-reported fatigue ratings on a 7-point scale before and after each block. A rating of 1 indicated no fatigue, 4 moderate fatigue, and 7 extreme fatigue. The average score represented the prevalent fatigue state for each block. Data samples—including VCT, EEG, and fNIRS features—were classified based on these fatigue ratings, using a threshold of 4 to distinguish between low fatigue (< 4) and high fatigue (≥ 4) (29, 51, 52). This measure was derived from subjective feelings frequently reported by participants in our pilot study and previous research. It was also based on the recognized interplay between brain states and stimulation effects; specifically, conditions like arousal and sleepiness, which correlate with fatigue, significantly influence responses to electrical stimulation (23, 24, 26, 36, 53). Note that in the histogram of Fig. 3F, the first five bars include values from the lower edge up to but excluding the upper edge, while the sixth bar includes values up to and including the upper edge. Consequently, we hypothesized state-dependent tRNS effects during the experiment— that is, tRNS effects would vary under different levels of fatigue.

### EEG Data Collection, Preprocessing, and Computation of Alpha/Beta Power

EEG signals were recorded using a 64-channel electrode system (Easycap) and an EEG amplifier (BrainAmp, Brain Products GmbH, Germany), following the international 10-10 system. Electrodes were positioned over occipital channels (O1, O2, PO3, PO4, POz), with FCz as the reference electrode. Impedances were kept below 10 kΩ. The raw EEG data were preprocessed using the EEGLAB toolbox (54), including resampling to 250 Hz and band-pass FIR filtering from 0.1 to 80 Hz. Data were then segmented into epochs from –1800 to 1500 ms relative to stimulus onset, followed by visual artifact removal.

To compute prestimulus alpha and beta power, we estimated the power spectral density (PSD) of prestimulus EEG signals from each occipital electrode to quantify power within five specific frequency bands: delta (0.5–4 Hz), theta (4–8 Hz), alpha (8–13 Hz), beta (13–30 Hz), and gamma (30–45 Hz). We limited the gamma band to 45 Hz to avoid contamination from 50 Hz line noise and potential artifacts at higher frequencies. The PSD was estimated using Welch’s method with a Hamming window length of 200 samples, 50% overlap (100 samples), and an FFT length of 256 points, balancing spectral resolution and variance reduction. We calculated the power in each frequency band by averaging the PSD over the respective frequency ranges and converted these values to decibels (dB) to facilitate comparison across bands and electrodes. All computations were performed using MATLAB R2023b with built-in functions (pwelch and bandpower), ensuring consistency and reproducibility.

### fNIRS Data Collection, Preprocessing, and Computation of HbO Amplitude

As EEG data cannot be collected during tRNS due to electrical artifacts (27), we measured cortical excitability during stimulation with fNIRS, which were collected using a NIRSport2 system (NIRx Medical Technologies, LLC, New York, NY) at a sampling rate of 10.1725 Hz, employing wavelengths of 760 nm and 850 nm. The setup included six source probes and six detector probes, inserted into holders on a cap arranged according to the international 10-5 system (sources at PPOz, PPO3, PO1h, POO3, POO4, PPO4; detectors at PPO1, POOz, PO2h, PPO2, POO8, POO7), covering the occipital lobe through a network of 21 channels (Fig. 2B). The source-detector distances varied around an average of 32.743 ± 8.510 mm, optimized to accommodate simultaneous EEG and tRNS setups. To minimize external light interference, a dark over-cap was used. Changes in light attenuation were analysed using the modified Beer– Lambert law to derive relative changes in the concentrations of oxyhemoglobin (HbO) and deoxyhemoglobin (HbR). The HbO and HbR signals were high-pass filtered at 0.015 Hz and low-pass filtered at 0.1 Hz to eliminate high-frequency physiological noise and low-frequency baseline drifts. An artifact rejection threshold was established at the mean concentration plus or minus two standard deviations. Data collected at the beginning and end of measurements were discarded if HbO levels exceeded this threshold, adjusting for movement artifacts. After preprocessing, the fNIRS data reflected the relative changes of HbO and HbR in micromoles (μMol). To assess global variations during the resting periods, we calculated the mean time series of relative changes across all channels to derive the global signal in the occipital lobe. This averaging process enhanced the signal-to-noise ratio (SNR) and mitigated potential effects arising from variable channel distances. The amplitude of the HbO signal was quantified using the standard deviation of this averaged time series (55).

### Statistical Analysis of tRNS Effects

To evaluate the potential causal role of prestimulus alpha and beta power on VCT, we first compared tRNS effects on HbO amplitude, prestimulus alpha and beta power, and VCT across the two stimulation conditions (sham and tRNS) using Bayesian linear mixed models, implemented via the Stan modeling language (56) and the brms package (57). The Bayesian approach was adopted for its benefits in resolving non-convergence problems and facilitating the fitting of maximal varying effect structures (41). We then examined whether different stimulation conditions influenced fatigue ratings, which could potentially confound the categorization of low/high fatigue (58). Finally, we analysed how fNIRS HbO amplitude, EEG prestimulus power in different frequency bands, and VCT varied across blocks in the different stimulation conditions under different fatigue levels (low/high). Baseline measurements from Block 1 were included in the models as covariates, and the sham condition at Block 1 always served as the reference level. The results were evaluated by calculating the posterior probability of the contrast difference, *δ*, between estimated parameters corresponding to tRNS and sham conditions. Models hypothesized that *δ* ≠ 0, and evidence supporting these hypotheses was considered compelling if the posterior probability (*Pr*) exceeded 97.5% and the 95% HPD interval did not include zero (59, 60). Four sampling chains were run for 3000 iterations, with the first 1000 as the warm-up phase. The default priors of the brms package were applied. The specific models employed were as follows:

(1) tRNS effects on HbO amplitude, prestimulus alpha and beta power, VCT, and fatigue ratings (Fig. 3) HbO amplitude, prestimulus alpha and beta power, VCT, and fatigue ratings were modelled as a function of baseline measurements, stimulation condition, block, and the interaction between stimulation and block, with individual participants as random effects. The model syntax was:

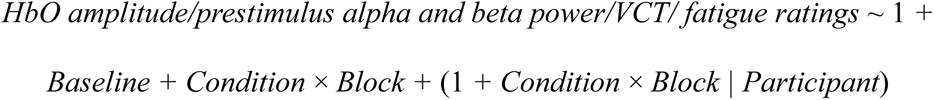
(2) State-dependent tRNS effects depending on fatigue level (Fig. 4&5): HbO amplitude, prestimulus power, and VCT were modeled to analyze the impact of fatigue level, the stimulation condition, block, and the interaction between stimulation and block. For the random effects part, we did not include the interaction of block due to increased complexity and implementation issues. The models were formulated as:

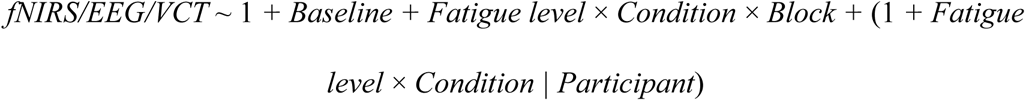

### Sensitivity analysis

To compare the contributions of EEG power in different frequency bands to VCT under varying fatigue states, we employed neural network models to fit the relationship between single-trial EEG power and VCT. After training, we computed the sensitivity matrix and performed eigenvalue decomposition to quantify feature importance. Finally, we conducted permutation tests to explore potential reasons why tRNS effects were observed only under low fatigue.

We chose neural network models to capture potential nonlinear relationships between EEG power and VCT. Two separate models were trained using data from the sham group under low and high fatigue conditions, respectively. Both models had identical architectures and hyperparameters: one hidden layer with 64 units, a batch size of 32, a maximum of 200 epochs, a learning rate of 0.001, an optimizer of Adam, and the ReLU activation function. The input *x_i_* consisted of 25 features representing EEG power across 5 frequency bands and 5 electrodes (denoted as *x*_*i*,1_, *x*_*i*,2_, …, *x*_*i*,25_, in Fig. 6A), and the output *y*_*i*_ was the single-trial VCT. Since VCT was measured every 40 trials within a block, each group of 40 samples shared the same VCT value. The datasets comprised 6,640 samples for low fatigue and 4,960 samples for high fatigue. To prevent overfitting, we split each dataset into training and validation subsets in an 8:2 ratio and applied early stopping with a patience of 5 epochs. Training for the low fatigue model stopped after 58 epochs (iteration 9628) and for the high fatigue model after 103 epochs (iteration 12782). Validation loss remained slightly lower than training loss in both cases, indicating model generalizability. The similar loss values between conditions suggest the models are comparable for cross-condition analysis. Models were implemented using the Machine Learning Toolbox in MATLAB R2023b.

After training the models, we computed the sensitivity matrix S for each fatigue state using the formula:

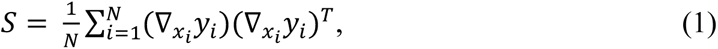

Where *N* is the number of samples, *y_i_* is the model’s output for the *i*-th sample, *x_i_* is the input vector for the i-th sample, and ∇_*x*_*i*__ *y_i_* is the gradient of the model’s output with respect to the input vector *x_i_*, i.e.,

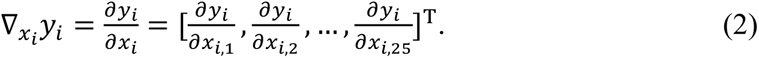

The sensitivity matrix captures how variations in input features affect the model’s output, providing a quantitative measure of feature importance and interactions. We then performed eigenvalue decomposition on *S*:

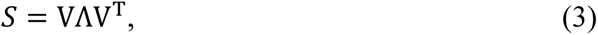

where V is a matrix, whose columns are the eigenvectors, and Λ is a diagonal matrix containing the corresponding eigenvalues. The eigenvalues were sorted in descending order. Larger eigenvalues correspond to directions in the input space where the output is more sensitive. For every eigenvalue, its corresponding eigenvector represents a specific “sensitivity direction”, with its components indicating the contribution of each feature along this direction. We focused on the eigenvectors associated with the largest eigenvalues (explaining 90% of the variance) and computed the mean absolute values of their components to quantify the sensitivity contributions of different frequency bands. We compared the contribution values for the alpha and beta bands (Fig. 6D, left panel).

To investigate why tRNS effects were observed only under low fatigue, we statistically compared the sensitivity contributions of different frequency bands between low and high fatigue states using permutation tests. We combined all samples from both fatigue states and randomly permuted their labels 10,000 times. In each permutation, we conducted the sensitivity analysis as described above to obtain the difference in contributions for each frequency band between low and high fatigue. We calculated *p*-values by determining the position of the observed difference within the distribution of permuted differences for each frequency band. False discovery rate (FDR) correction was applied for multiple comparisons.

To justify the hyperparameter combination, we optimized our neural network models by systematically varying key hyperparameters: the number of hidden units (32 and 64), batch sizes (32 and 64), and learning rates (0.01 and 0.001), resulting in eight different combinations. Early stopping with a patience of 5 epochs was employed to prevent overfitting. We selected these hyperparameters because they directly affect model capacity, training dynamics, and the ability to capture nonlinear relationships. We compared the validation loss across all combinations and selected the hyperparameter configuration that yielded the lowest validation loss for the final model due to its superior fit and generalization performance. This approach ensured that our model achieved optimal performance while maintaining robustness against overfitting. By systematically varying these hyperparameters and evaluating model performance based on validation loss, we demonstrated that our findings reflect intrinsic properties of the data rather than being artifacts of specific hyperparameter settings or model characteristics. The comparison results can be found in Table S13 and Fig. S2.

## Funding

This work was supported in part by [the National Natural Science Foundation of China] (Nos. 82272114, 32361143787), and in part by [Shenzhen-Hong Kong Institute of Brain Science-Shenzhen Fundamental Research Institutions] (2022SHIBS0003).

## Data and code availability

The data and code will be available in the open science framework at the time of publication.

## Conflict of interest

The authors have no relevant financial or non-financial interests to disclose.

## Classification

Biological Sciences: Neuroscience

## Notes

### Competing Interest Statement

The authors have declared no competing interest.

