## Supplemental Information for "Modulating Prestimulus Alpha and Beta Power with tRNS Establishes Their Causal Role in Visual Perception"

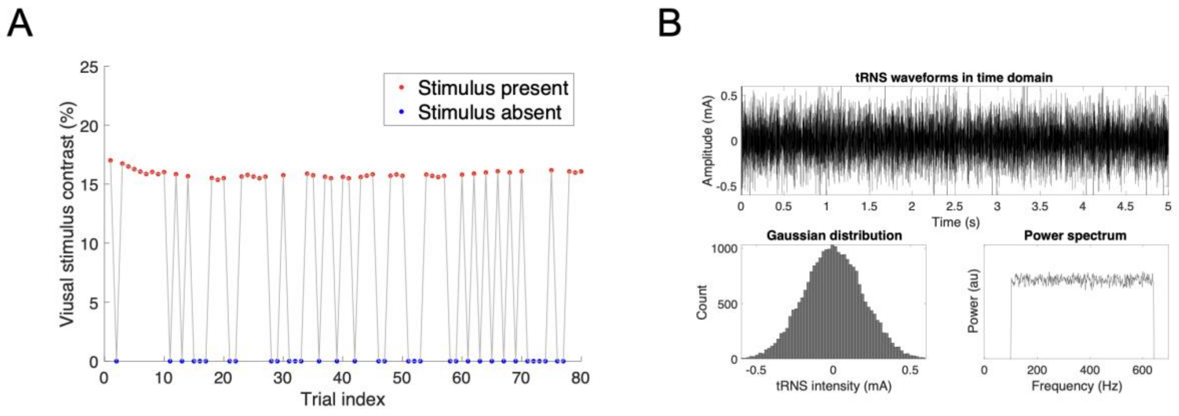


**Fig. S1.** (*A*) Within each block, visual stimuli were presented in 60% of trials (denoted by red dots), while stimuli were absent in the remaining trials (denoted by blue dots). (*B*) The tRNS current waveform had intensities normally distributed, with 99% of values within the peak-to-peak amplitude of 1 mA. The power spectrum illustrates the high-frequency band (100–640 Hz) tRNS used in the study.

**Table S1.** Effects of tRNS on fNIRS HbO amplitude (refer to Fig. 3A).

This table reported the contrast results of fNIRS HbO amplitude between sham and tRNS conditions obtained from Bayesian linear mixed model. It includes the estimated contrast coefficients (*Est*), the lower and upper ranges of the 95% highest probability density (Lower *HPD*: *LH*; Upper *HPD*: *UH*) for these estimates, and the posterior probability (*Pr*) indicating the presence of contrast difference. The hypotheses stating that contrast difference existed were formulated. The evidence was considered compelling to support the hypothesis if the posterior probability exceeds 97.5% and 95% *HPD* does not include 0, as shown by bold line. For the associated linear mixed models, refer to the Methods section.

| Contrast | Block | *Est* | *LH* | *UH* | *Pr* |
|---|---|---|---|---|---|
| Sham - tRNS | 2 | –0.079 | –0.183 | 0.029 | 0.929 |
| Sham - tRNS | 3 | –0.109 | –0.209 | 0.004 | 0.979 |
| Sham - tRNS | 4 | –0.094 | –0.194 | 0.019 | 0.958 |
| **Sham - tRNS** | **5** | **–0.169** | **–0.278** | **–0.063** | **0.999** |

**Table S2.** Effects of tRNS on alpha power (refer to Fig. 3B).

| Contrast | Block | *Est* | *LH* | *UH* | *Pr* |
|---|---|---|---|---|---|
| Sham - tRNS | 2 | −0.257 | −0.727 | 0.203 | 0.853 |
| Sham - tRNS | 3 | −0.144 | −0.629 | 0.307 | 0.730 |
| Sham - tRNS | 4 | −0.054 | −0.509 | 0.436 | 0.589 |
| Sham - tRNS | 5 | 0.093 | −0.396 | 0.566 | 0.350 |

**Table S3.** Effects of tRNS on beta power (refer to Fig. 3C).

| Contrast | Block | *Est* | *LH* | *UH* | *Pr* |
|---|---|---|---|---|---|
| Sham - tRNS | 2 | −0.147 | −0.516 | 0.217 | 0.799 |
| Sham - tRNS | 3 | −0.044 | −0.386 | 0.333 | 0.594 |
| Sham - tRNS | 4 | 0.078 | −0.291 | 0.431 | 0.670 |
| Sham - tRNS | 5 | 0.142 | −0.203 | 0.523 | 0.780 |

**Table S4.** Effects of tRNS on VCT (refer to Fig. 3D).

| Contrast | Block | *Est* | *LH* | *UH* | *Pr* |
|---|---|---|---|---|---|
| Sham - tRNS | 2 | 0.042 | −0.229 | 0.324 | 0.623 |
| Sham - tRNS | 3 | 0.181 | −0.079 | 0.459 | 0.913 |
| Sham - tRNS | 4 | 0.207 | −0.069 | 0.472 | 0.944 |
| Sham - tRNS | 5 | 0.184 | −0.085 | 0.448 | 0.922 |

**Table S5.** Effects of tRNS on fatigue ratings (refer to Fig. 3E).

| Contrast | Block | *Est* | *LH* | *UH* | *Pr* |
|---|---|---|---|---|---|
| Sham - tRNS | 2 | −0.034 | −0.234 | 0.141 | 0.639 |
| Sham - tRNS | 3 | −0.008 | −0.214 | 0.224 | 0.527 |
| Sham - tRNS | 4 | 0.186 | −0.209 | 0.598 | 0.824 |
| Sham - tRNS | 5 | 0.088 | −0.319 | 0.518 | 0.672 |

**Table S6.** Effects of tRNS on fNIRS HbO amplitude under different fatigue states (refer to Fig. 4A). This table reported the contrast results of fNIRS HbO amplitude under low and high fatigue states between sham and tRNS conditions obtained from Bayesian linear mixed model.

| | | HbO: Low fatigue | | | | HbO: High fatigue | | | |
|---|---|---|---|---|---|---|---|---|---|
| Contrast | Block | *Est* | *LH* | *UH* | *Pr* | *Est* | *LH* | *UH* | *Pr* |
| Sham - tRNS | 2 | –0.096 | –0.230 | 0.028 | 0.930 | –0.035 | –0.197 | 0.113 | 0.673 |
| Sham - tRNS | 3 | –0.133 | –0.276 | –0.005 | 0.974 | –0.071 | –0.210 | 0.072 | 0.839 |
| Sham - tRNS | 4 | –0.128 | –0.263 | 0.001 | 0.969 | –0.058 | 0.206 | 0.088 | 0.781 |
| **Sham - tRNS** | **5** | **–0.276** | **–0.405** | **–0.137** | **1** | –0.039 | –0.183 | 0.109 | 0.706 |

**Table S7.** Effects of tRNS on **alpha** power under different fatigue states (refer to Fig. 4B). This table reported the contrast results of alpha power under low and high fatigue states between sham and tRNS conditions obtained from Bayesian linear mixed model.

| | | Alpha power: Low fatigue | | | | Alpha power: High fatigue | | | |
|---|---|---|---|---|---|---|---|---|---|
| Contrast | Block | *Est* | *LH* | *UH* | *Pr* | *Est* | *LH* | *UH* | *Pr* |
| Sham - tRNS | 2 | –0.354 | –0.897 | 0.195 | 0.900 | –0.124 | –0.744 | 0.542 | 0.650 |
| Sham - tRNS | 3 | –0.129 | –0.645 | 0.423 | 0.677 | –0.248 | –0.899 | 0.368 | 0.786 |
| Sham - tRNS | 4 | 0.330 | –0.224 | 0.873 | 0.882 | –0.633 | –0.253 | 0.032 | 0.972 |
| **Sham - tRNS** | **5** | **0.582** | **0.034** | **1.162** | **0.977** | –0.453 | –1.013 | 0.156 | 0.930 |

**Table S8.** Effects of tRNS on **beta** power under different fatigue states (refer to Fig. 4C). This table reported the contrast results of beta power under low and high fatigue states between sham and tRNS conditions obtained from Bayesian linear mixed model.

| | | Beta power: Low fatigue | | | | Beta power: High fatigue | | | |
|---|---|---|---|---|---|---|---|---|---|
| Contrast | Block | *Est* | *LH* | *UH* | *Pr* | *Est* | *LH* | *UH* | *Pr* |
| Sham - tRNS | 2 | –0.107 | –0.510 | 0.307 | 0.698 | –0.229 | –0.809 | 0.326 | 0.819 |
| Sham - tRNS | 3 | –0.013 | –0.433 | 0.441 | 0.477 | –0.080 | –0.628 | 0.475 | 0.621 |
| Sham - tRNS | 4 | 0.297 | –0.112 | 0.725 | 0.919 | –0.245 | –0.798 | 0.333 | 0.800 |
| **Sham - tRNS** | **5** | **0.482** | **0.047** | **0.927** | **0.983** | –0.209 | –0.755 | 0.330 | 0.782 |

**Table S9.** Effects of tRNS on **VCT** under different fatigue states (refer to Fig. 4C).
This table reported the contrast results of VCT under low and high fatigue states between sham and tRNS conditions obtained from Bayesian linear mixed model.

| | | VCT: Low fatigue | | | | VCT: High fatigue | | | |
|---|---|---|---|---|---|---|---|---|---|
| Contrast | Block | *Est* | *LH* | *UH* | *Pr* | *Est* | *LH* | *UH* | *Pr* |
| Sham - tRNS | 2 | 0.015 | –0.273 | 0.309 | 0.546 | 0.053 | –0.308 | 0.402 | 0.617 |
| Sham - tRNS | 3 | 0.163 | –0.116 | 0.462 | 0.862 | 0.229 | –0.119 | 0.581 | 0.900 |
| Sham - tRNS | 4 | 0.245 | –0.049 | 0.551 | 0.952 | 0.206 | –0.154 | 0.556 | 0.871 |
| **Sham - tRNS** | **5** | **0.307** | **0.001** | **0.603** | **0.978** | 0.092 | –0.258 | 0.414 | 0.702 |

**Table S10.** Effects of tRNS on **delta power** under different fatigue states (refer to Fig. 5A).
This table reported the contrast results of delta power under low and high fatigue states between sham and tRNS conditions obtained from Bayesian linear mixed model.

| | | Delta power: Low fatigue | | | | Delta power: High fatigue | | | |
|---|---|---|---|---|---|---|---|---|---|
| Contrast | Block | *Est* | *LH* | *UH* | *Pr* | *Est* | *LH* | *UH* | *Pr* |
| Sham - tRNS | 2 | –0.542 | –2.396 | 1.616 | 0.697 | –0.574 | –3.256 | 2.082 | 0.665 |
| Sham - tRNS | 3 | –0.322 | –2.312 | 1.742 | 0.621 | 0.608 | –2.189 | 3.153 | 0.675 |
| Sham - tRNS | 4 | 0.554 | –1.529 | 2.648 | 0.700 | 0.286 | –2.492 | 2.899 | 0.584 |
| Sham - tRNS | 5 | 1.047 | –1.025 | 3.206 | 0.838 | 0.825 | –1.760 | 3.232 | 0.746 |

**Table S11.** Effects of tRNS on **theta power** under different fatigue states (refer to Fig. 5B).
This table reported the contrast results of delta power under low and high fatigue states between sham and tRNS conditions obtained from Bayesian linear mixed model.

| | | Theta power: Low fatigue | | | | Theta power: High fatigue | | | |
|---|---|---|---|---|---|---|---|---|---|
| Contrast | Block | *Est* | *LH* | *UH* | *Pr* | *Est* | *LH* | *UH* | *Pr* |
| Sham - tRNS | 2 | –0.291 | –0.829 | 0.234 | 0.858 | –0.058 | –0.735 | 0.593 | 0.568 |
| Sham - tRNS | 3 | –0.405 | –0.982 | 0.098 | 0.932 | 0.042 | –0.607 | 0.683 | 0.548 |
| Sham - tRNS | 4 | 0.136 | –0.366 | 0.685 | 0.701 | –0.326 | –0.970 | 0.350 | 0.839 |
| Sham - tRNS | 5 | 0.142 | –0.383 | 0.708 | 0.691 | –0.242 | –0.854 | 0.355 | 0.777 |

**Table S12.** Effects of tRNS on **gamma power** under different fatigue states (refer to Fig. 5C). This table reported the contrast results of gamma power under low and high fatigue states between sham and tRNS conditions obtained from Bayesian linear mixed model.

| | | Gamma power: Low fatigue | | | | Gamma power: High fatigue | | | |
|---|---|---|---|---|---|---|---|---|---|
| Contrast | Block | *Est* | *LH* | *UH* | *Pr* | *Est* | *LH* | *UH* | *Pr* |
| Sham - tRNS | 2 | 0.057 | −0.549 | 0.678 | 0.572 | −0.363 | −1.285 | 0.582 | 0.781 |
| Sham - tRNS | 3 | 0.059 | −0.547 | 0.718 | 0.572 | −0.015 | −0.889 | 0.924 | 0.512 |
| Sham - tRNS | 4 | 0.120 | −0.490 | 0.766 | 0.644 | 0.050 | −0.914 | 0.982 | 0.541 |
| Sham - tRNS | 5 | 0.586 | −0.076 | 1.271 | 0.961 | −0.325 | −1.261 | 0.498 | 0.772 |

### Hyperparameter Selection in Sensitivity Analysis

To ensure the robustness and reliability of our neural network models, we conducted a systematic exploration of key hyperparameters. Specifically, we varied the number of hidden units (32 and 64), batch sizes (32 and 64), and learning rates (0.01 and 0.001), resulting in a total of eight combinations. The early stopping patience was consistently set to 5 epochs across all experiments.

The selection of 32 and 64 hidden units was based on balancing model complexity with the risk of overfitting, considering our input size of 25 features. Choosing hidden layer sizes that are not excessively larger than the input size helps prevent over-parameterization and ensures better generalization to unseen data (1). This aligns with established guidelines suggesting that the number of hidden units should be proportional to the input size to maintain computational efficiency and training stability (2). Larger hidden layer sizes, such as 128 units, were avoided to reduce the risk of overfitting given our dataset size and to maintain computational efficiency (3).

Batch sizes of 32 and 64 were chosen to balance computational efficiency and convergence speed. Smaller batch sizes can lead to noisy gradient estimates, while larger batch sizes may require more memory and could potentially slow down convergence (4). By selecting these batch sizes, we aimed to achieve efficient training while maintaining the quality of gradient updates.

We experimented with learning rates of 0.01 and 0.001 to explore the impact of different optimization speeds on model performance. A higher learning rate like 0.01 allows for faster convergence but may risk overshooting minima, while a lower learning rate like 0.001 provides more stable convergence at the expense of longer training times (5). Exploring both values enabled us to assess the sensitivity of our models to this critical hyperparameter.

The early stopping patience was set to 5 epochs based on recommendations from the literature and standard practices observed in machine learning frameworks. This choice balances the need for sufficient training time to allow the model to improve with the goal of preventing overfitting and ensuring efficient use of computational resources (6, 7). Setting the patience to 5 prevents the early stopping mechanism from being too sensitive to minor fluctuations in validation loss, allowing the training process to focus on longer-term trends in model improvement.

We evaluated each hyperparameter combination by comparing the validation loss obtained during training (Table S13). The combination that resulted in the lowest validation loss was selected for the final model due to its better fit and generalization performance. This approach allowed us to identify the most effective hyperparameter configuration for our data, ensuring that the model achieves optimal performance while avoiding overfitting. We found that when batch size equals 32 or 64 and other hyperparameters were fixed, the validation losses were quite comparable (Table S13). Therefore, we also tried the batch size of 64 and performed the same procedures for sensitivity analysis (Fig. S2). The results were consistent with Fig. 6.

By systematically varying these hyperparameters and evaluating model performance based on validation loss, we ensured that our findings are robust and not artifacts of specific hyperparameter choices.

**Table S13.** Comparison of validation loss across different hyperparameter combinations. This table reported the validation losses by using different hyperparameter combinations in neural network modelling of the sensitivity analysis. The bold rows yielded comparable results.

| Hidden size | Batch size | Learning rate | Validation loss |
|---|---|---|---|
| 32 | 32 | 0.01 | Low: 0.155; high: 0.167 |
| 32 | 64 | 0.01 | Low: 0.153; high: 0.177 |
| 32 | 32 | 0.001 | Low: 0.139; high: 0.162 |
| 32 | 64 | 0.001 | Low: 0.140; high: 0.159 |
| 64 | 32 | 0.01 | Low: 0.152; high: 0.167 |
| 64 | 64 | 0.01 | Low: 0.135; high: 0.164 |
| **64** | **32** | **0.001** | **Low: 0.130; high: 0.143** |
| **64** | **64** | **0.001** | **Low: 0.125; high: 0.150** |

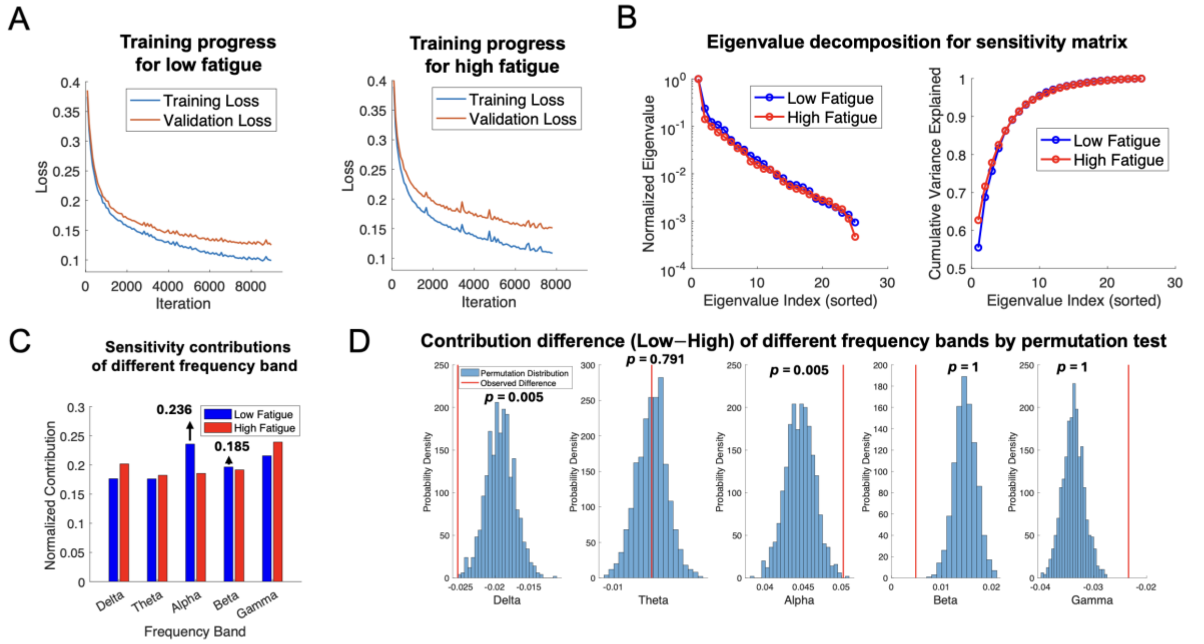


**Fig. S2.** Sensitivity analysis for another hyperparameter combination candidate: hidden size = 64, batch size = 64, learning rate = 0.001, patience = 5. (*A*) To prevent overfitting, samples under each fatigue state were divided into training and validation subsets. Early stopping was employed during model training. (*B*)

Eigenvalue decomposition of the sensitivity matrix was performed. Eigenvalues explaining 90% of the variance were selected to calculate the sensitivity contributions of different frequency bands to VCT. (*C*) Under low fatigue, the alpha band showed a larger sensitivity contribution to VCT than the beta band. (*D*) The alpha band had higher sensitivity under low fatigue compared to high fatigue, whereas the beta band did not show a significant difference between fatigue states. Note that the $p$ values was FDR corrected.
